# MCL1 may not mediate chemoresistance

**DOI:** 10.1101/2025.08.22.671824

**Authors:** Kylin A. Emhoff, Kunho Chung, Dongmei Zhang, Belinda Willard, Timothy Chan, Babal Kant Jha, Shaun R. Stauffer, Jesse A. Coker, Jan Joseph Melenhorst

## Abstract

The anti-apoptotic BCL2 family member MCL1 is overexpressed in many cancers and has been linked to chemoresistance. Unlike other BCL2 family members, MCL1 displays both well-defined mitochondrial anti-apoptotic activities and also emerging nuclear functions. Prior reports suggest that MCL1 enters the nucleus during chemotherapy and promotes chemoresistance by influencing cell cycle progression and DNA repair. These nuclear roles of MCL1, however, remain poorly characterized. Using a newly validated monoclonal antibody across several cell lines and treatments, we find no evidence that MCL1 enhances chemoresistance or preferentially accumulates in the nucleus after drug exposure. Proximity biotinylation identified novel nuclear MCL1 interactors but did not recover previously reported DNA repair or cell cycle partners. Thus, while MCL1 does reach the nucleus and interact with nuclear proteins, our data do not support a role for MCL1 in chemoresistance. Further work is needed to clarify the functional significance of nuclear MCL1.

**SIGNFIGANCE:** Previous studies have implicated MCL1 in promoting chemoresistance via interactions with DNA repair machinery in the nucleus. Using a validated, monoclonal anti-MCL1 antibody, we were unable to replicate these data. We report that MCL1 neither confers chemoresistance, translocates to the nucleus during chemotherapy treatment, nor interacts with DNA repair proteins in live cancer cells.

## INTRODUCTION

The upregulation of anti-apoptotic B-cell leukemia/lymphoma 2 (BCL2) family members is a major mechanism of chemoresistance in many cancers.^1^ These proteins sequester mitochondrial apoptosis inducers such as BAX, BAD, and BIM through BCL2-homology (BH) domain-mediated protein-protein interactions (PPIs).^2^ Myeloid Cell Leukemia 1 (MCL1) is a key BCL2 family member that binds to BAK, BIM, BID, and PUMA,^3^ is overexpressed in several malignancies, including cervical and colorectal cancers,^4^ and correlates with poor prognosis and therapeutic failure.^5^ Tumors with high MCL1 levels quickly acquire resistance to small molecule inhibitors of BCL2 and/or BCL-XL because MCL1 compensates for their inhibition.^6–8^ These observations prompted the development of potent, selective MCL1 inhibitors that compete with pro-apoptotic BH3 peptides at MCL1’s BH3 binding groove and trigger cell death.^9–13^ However, these inhibitors have shown on-target cardiotoxicity and a narrow therapeutic window in clinical studies.^14^ Safer strategies to curb MCL1’s oncogenic activity are therefore needed.

MCL1 is a 37 kDa protein with three C-terminal BH domains and an extended, disordered N-terminus containing PEST (Pro, Glu, Ser, Thr) motifs. Phosphorylation of the PEST domains by glycogen synthase kinase-3 beta (GSK3β), c-Jun N-terminal kinases (JNKs), extracellular signal-regulated kinases (ERKs), and cyclin-dependent kinases (CDKs)^15^ control MCL1’s short half-life (0.5 - 4 h).^3, 16–18^ MCL1 knock-out (KO) is embryonically lethal, which is unique relative to other BCL2 family members, underscoring specialized functions of MCL1 beyond apoptosis regulation.^19^ Reported non-canonical activities include roles in embryonic implantation,^20^ lymphocyte maintenance and hematopoiesis,^21–23^ mitochondrial homeostasis,^24, 25^ fatty acid oxidation,^26–28^ and autophagy.^29–31^

Many studies have linked these non-canonical functions to nuclear localized MCL1, which is usually tethered to the mitochondria via a C-terminal helix.^32^ MCL1, including especially a C-terminally truncated short nuclear form (snMCL1), does reach the nucleus.^33^ MCL1 interacts with the radiation-inducible immediate-early gene (IEX-1) for nuclear import.^34^ Additionally, nuclear MCL1 is cell cycle regulated and peaks during the S/G2 phase.^35^ Nuclear accumulation of MCL1 has been reported after irradiation or treatment with hydroxyurea, etoposide, oxaliplatin, or doxorubicin,^34, 36–38^ suggesting a role for nuclear MCL1 in DNA damage response. MCL1 has been observed at γH2AX-positive sites and linked to chemotherapy-induced senescence.^39–42^ It has been suggested that nuclear MCL1 slows proliferation by arresting cells at S and G2/M checkpoints: MCL1 inhibits PCNA^43^ and CDK1^33^ while activating CHK1,^36^ thereby favoring DNA repair via homologous recombination (HR) over non-homologous end joining (NHEJ).^35, 44^ However, other studies implicate MCL1 as a pro-proliferative protein.^45–47^ Importantly, many of these studies characterizing nuclear MCL1 relied on a now discontinued anti-MCL1 polyclonal antibody (SCBT #sc-819).^33–36, 39, 40^

Because nuclear MCL1 is implicated in proliferation and chemoresistance, we hypothesized that disrupting its nuclear interactions might offer a broader therapeutic window than BH3 blockade. Previous work relied on the discontinued antibody SCBT #sc-819, requiring us to validate a distinct monoclonal antibody with confirmed specificity. Using this reagent, we detected a stable nuclear pool of MCL1 across cell lines but saw no chemotherapy-induced nuclear translocation or MCL1-dependent chemoresistance. Proximity biotinylation identified >12 novel nuclear interactors of MCL1, none related to DNA damage or cell cycle control previously reported by co- immunoprecipitation (co-IP). Collectively, our findings do not support the model that chemotherapy drives MCL1 into the nucleus to mediate chemoresistance.

## MATERIALS AND METHODS

### Cell lines

Wild-type (WT) cell lines HCT116, HeLa, HEK293, and mouse embryonic fibroblasts (MEFs) were obtained from ATCC through the Cell Services Core Facility, Cleveland Clinic Foundation. HCT116^MCL1-KO^ and HeLa^MCL1-KO^ cell lines were custom generated by Ubigene Biosciences (Guangzhou Science City, Guangdong province, China); HEK293^MCL1-KO^ cells were custom generated by Abcam Inc (Waltham, MA). HeLa, HEK293, and MEFs were cultured in high glucose Dulbecco’s Modified Eagle Medium (DMEM) supplemented with L-glutamine and sodium pyruvate and 10% (v/v) fetal bovine serum (FBS). HCT116 cells were cultured in McCoy’s 5A medium supplemented with L-glutamine and 10% (v/v) FBS. All cell lines were maintained at 37°C, 5% CO2.

### Transfections

WT and MCL1^KO^ cell lines were plated in 6-well plates at 0.3 x 10^6^ cells/well for ≥ 16 h. Except for MEFs, cells were transfected with MCL1^WT^, MCL1^NLS^, GFP^TurboID^ or MCL1^TurboID^ plasmid constructs (Twist Biosciences, South San Francisco, CA) using Lipofectamine 2000 according to the manufacturer’s protocol (#11668027, ThermoFisher). Transfections took place for 48 h. MEFs were transfected with a *Silencer*® Select scramble (SCR.) siRNA or a *Silencer*® Select MCL1 siRNA (Life Technologies-Invitrogen, Carlsbad, CA) for 24 h using Lipofectamine RNAiMAX according to the manufacturer’s protocol (#13778075, ThermoFisher).

### Subcellular fractionation & western blot

Cell pellets (1.0 x 10^6^ cells per sample) were first resuspended in 25 µL cytosolic lysis buffer (10 mM HEPES pH 8.0, 10 mM KCl, 1.5 mM MgCl2, 0.34 M sucrose, 10% glycerol, 1 mM DTT, 1X protease inhibitor cocktail) and 0.1% Triton X-100. Resuspended pellets were incubated on ice for 5 min before centrifugation for 4 min at 4°C, 1300 x g. Supernatants containing cytosolic proteins were transferred to fresh microcentrifuge tubes and then centrifuged at 4°C, max speed (17200 x g) for 10 min. Remaining pellets were washed with 50 µL of cytosolic lysis buffer by centrifuging for 4 min at 1300 x g. Pellets were then resuspended in 25 µL chromatin extraction buffer (1X D-PBS and 5X benzonase). Resuspended pellets were incubated at 37°C for 25 min and then sonicated in a water bath at room temperature for 5 min. 10 µL of 2X loading dye was added to each sample and boiled at 95.5°C for 10 min.

Samples were loaded onto gradient gels that were run using 1X running buffer (25 mM Tris, 192 mM glycine, 0.1% SDS, pH 8.3). Proteins were transferred from gels onto nitrocellulose membranes (0.2 µm pore size) using transfer buffer (25 mM Tris, 192 mM glycine, 20% methanol, pH 8.0). Blots were blocked with 5% BSA in TBST for 1 h at room temperature. Blots were then incubated with MCL1 (1:1000 dilution, anti-rabbit; #5453, Cell Signaling Technology), FLAG (1:1000 dilution, anti-rabbit; #14793S, Cell Signaling Technology), GAPDH (1:1000 dilution, anti-mouse; #sc-47724, Santa Cruz Biotechnology), β-ACTN (1:1000 dilution, anti-mouse; #3700, Cell Signaling Technology), HISTONE H3 (1:2000 dilution, anti-rabbit; #4499, Cell Signaling Technology) or WDR5 (1:1000 dilution, anti-rabbit; #13105, Cell Signaling Technology) antibodies overnight at 4°C. The next day blots were incubated with secondary antibodies (anti-rabbit IgG HRP-linked; #AP307P, EMD Millipore Corp; anti-mouse IgG HRP-linked; #170-6516, Bio-Rad) for 1.5 h at room temperature. Enhanced chemiluminescent substrate solution (#32106, Thermo Scientific) was added to the blots and chemiluminescent signals were obtained on the Odyssey® M imager (LICORbio™, Lincoln, NE).

### Immunofluorescence (IF) microscopy

Cells were washed with cold 1X D-PBS and fixed with 4% paraformaldehyde for 15 min on ice. Cells were then permeabilizing with 0.3% Triton X-100 for 10 min on ice. Cells were then blocked with 5% BSA in PBST for 1 h at room temperature. Simultaneous primary antibody incubation then took place in which cells were incubated with an MCL1 antibody (1:1400 dilution, anti-rabbit; #5453, Cell Signaling Technology) and a TOM20 antibody (1:500 dilution, anti-mouse; #sc-17764, Santa Cruz Biotechnology) overnight at 4°C. The next day cells were simultaneously incubated with Alexa Fluor 488 (1:1000 dilution, anti-rabbit; #4412, Cell Signaling Technology) and Alexa Fluor 594 (1:1000 dilution, anti-mouse; A-11005, Invitrogen) secondary antibodies for 1 h in the dark at room temperature. Coverslips were mounted onto microscope slides with DAPI-containing mounting media. IF images were obtained using 40x magnification on a Leica DM5500B upright microscope equipped for fluorescence (Leica Microsystems, GmbH, Wetzlar, Germany) in our Imaging Core Facility, Cleveland Clinic Foundation.

### Cell viability assays

WT and MCL1^KO/KD^ cell lines were plated in 96-well plates at 5.0 x 10^3^ cells/well for ≥ 16 h. HCT116^MCL1-KO^ and HeLa^MCL1-KO^ cells were first transfected with MCL1^WT^ or MCL1^NLS^ plasmid constructs before being plated in 96-well plates. Moreover, MEFs were first transfected with SCR. siRNA or MCL1 siRNA. The next day cells were treated with the indicated concentrations of paclitaxel (PACL), vincristine (VINC), vinorelbine (VINO), doxorubicin (DOX), or etoposide (ETOPO) (MedChemExpress) for 48 h. CellTiter-Glo cell viability assay (CTG®; #G7573, Promega) was performed after 48-h compound treatment, and luminescence (LU) readings were obtained on a BioTek Cytation 5 microplate reader (Agilent, Santa Clara, CA). For the live-cell imaging experiments, CTG was performed after 7 days in the Incucyte®.

All cell viability data was analyzed in GraphPad Prism by normalizing LU values to % viability and graphing average % viability vs. log[compound] (M). Positive control implemented for these experiments was 1 µM staurosporine (STS), whereas vehicle control was 0.2% DMSO. The average LU value from the 0.2% DMSO vehicle control samples was set to “100% viability”, whereas the average luminescence value from the 1 μM STS samples was set to “0% viability”.

### Live-cell imaging

20 μL of media was added into each tested well of 384-well plates. Plates were then stamped with 50 nL of indicated concentrations of PACL, VINC, DOX or ETOPO (MedChemExpress) using a Labcyte Echo® 550 acoustic liquid handler (Beckman Coulter Life Sciences, Indianapolis, IN). After plate stamping, WT and KO cell lines were plated at 500 cells/well. Plates were then placed in the Incucyte® SX5 (Sartorius, Göttingen, Germany) for 7 days to monitor % phase object confluence. Area-under-the-curve (AUC) values were obtained for each % confluence vs. time growth curve and heatmaps were generated in GraphPad Prism.

### Nuclear translocation assays

HCT116 and HeLa cells were plated in 6-well plates at 0.3 x 10^6^ cells/well. The following day, cells were treated with 0.2 μM doxorubicin, 15 μM etoposide, 10 μM paclitaxel, 10 μM oxaliplatin, 2 μM 5-fluorouracil, or 0.1 μg/mL colcemid overnight (20 h) or with 0.15 μM, 1.5 μM, or 15 μM etoposide for 3 h. Subcellular fractionation western blot and IF staining were then performed.

### Proximity biotinylation & peptide preparation

Culture media from HCT116 and HeLa cells transfected with GFP^TurboID^ or MCL1^TurboID^ was removed and replaced with 67 µM biotin. Cells were incubated with exogenous biotin for 15-20 min at 37°C. Cells were then washed four times with ice-cold 1X D-PBS and then harvested with trypsin. Cell pellets were resuspended in 1 mL complete lysis buffer (RIPA lysis buffer, Thermo Scientific, catalog #89901; 1X protease inhibitor cocktail and 1X benzonase) and incubated on ice for 30 min. Resuspensions were centrifuged at max speed (17200 x g) for 15 min. at 4°C and supernatants were transferred to fresh microcentrifuge tubes.^48^ Protein concentration for each sample was estimated using BCA assay.

Biotinylated proteins in each sample were captured using streptavidin magnetic beads (#88817, Thermo Scientific) on a rotator overnight at 4°C. The following day, beads were collected, and supernatants ("flow-through" samples) were transferred to fresh microcentrifuge tubes and stored at -20°C. Beads were washed according to Cho et al.^48^

Protein elution was performed by on-bead digest using a solution consisting of 2 M urea in 50 mM Tris-HCl (pH 7.5), 1 mM DTT and 0.2 µg trypsin (#V5113, Promega) for 1 h at room temperature. Beads were collected, and eluates were transferred to fresh Protein LoBind microcentrifuge tubes. Beads were washed twice with 2 M urea in 50 mM Tris-HCl (pH 7.5); these washes were combined with the eluates. Average protein concentration for each eluate was estimated by measuring absorbance at 280 nm on a NanoDrop ND-1000 spectrophotometer (Thermo Scientific, Waltham, MA). DTT was added to a final concentration of 4 mM and incubated for 30 min at room temperature. Iodoacetamide was added to a final concentration of 10 mM and incubated for 1 h at room temperature in the dark. 0.25 µg trypsin was added to the eluates and digested overnight at 37°C.^48^

After overnight trypsin digestion, eluates were acidified for desalting by adding trifluoroacetic acid (TFA) to a final concentration of 1% (v/v). Using C18 OMIX desalting tips (#87784, Thermo Scientific), each eluate sample was desalted. Briefly, tip was washed three times with 100 µL 50% acetonitrile (ACN); tip was then equilibrated three times with 100 µL 0.1% TFA. Peptides from eluate of one sample were then loaded onto the tip with the "flow-through" solution being transferred to a fresh microcentrifuge tube. After loading peptides onto the C18 column, tip was rinsed five times with 100 µL wash buffer (0.1% TFA and 5% ACN). Peptides were slowly eluted with elution buffer (0.1% formic acid (FA) and 60% ACN) into a Protein LoBind tube. Desalting steps were repeated for the rest of the samples. Eluted peptide samples were dried using a SpeedVac lyophilizer.

### Liquid chromatography and tandem mass spectrometry

Mass spectrometry analysis of the peptide samples was performed by our Proteomics & Metabolomics Core Facility, Cleveland Clinic Foundation. An Orbitrap Exploris 480 mass spectrometer equipped with a Vanquish Neo uHPLC system (Thermo Fisher Scientific, Waltham, MA) was used to acquire mass spectra. The HPLC column was a 50 cm x 75 µm id Easy Spray Pepmap NEO C18, 2μm, 100 Å reversed- phase capillary chromatography column. 5 μL volumes of the extracts were injected, and the peptides were eluted from the column by an ACN/0.1% FA gradient at a flow rate of 0.3 μL/min and were introduced into the source of the mass spectrometer on-line. Digests were analyzed using the data-dependent multitask capability of the instrument, acquiring full-scan mass spectra to determine peptide molecular weights and product ion spectra to determine amino acid sequence in successive instrument scans. The Exploris data was analyzed by using all HCD spectra collected in the experiment to search the human SwissProt database using Proteome Discoverer 2.5. The SAINT (**<u>S</u>**ignificance **<u>A</u>**nalysis of **<u>Int</u>**eractome) algorithm was used to process mass spectrometry data and identify specific interactors of MCL1.^49^

### Statistical analysis

pGI50 values were compared by one-way ANOVAs with multiple comparisons, which were performed in GraphPad Prism. Multiplicity adjusted P values were obtained for each comparison to determine statistical significance; P > 0.05 (ns), P ≤ 0.05 (*), P ≤ 0.01 (**); ns = not significant.

### Data availability

All data is available within the main text and its supplemental information and files.

## RESULTS

### Validation of an anti-MCL1 monoclonal antibody

We first validated the monoclonal anti-MCL1 antibody CST #5453 as a substitute for the discontinued polyclonal antibody SCBT #sc-819.^33–36, 39, 40^ Western blots with MCL1^KO^ HCT116, HeLa, and HEK293 human lines, as well as MCL1^KD^ MEFs, confirmed specificity for human and mouse MCL1 (**Fig. 1A-D**). IF gave the same results (**Fig. 1E**). These experiments also validated the MCL1^KO^ or MCL1^KD^ status of all cell lines. MEFs were analyzed by siRNA-mediated MCL1^KD^ rather than KO because their limited proliferation hampers clonal isolation.^50^

**Figure 1.**
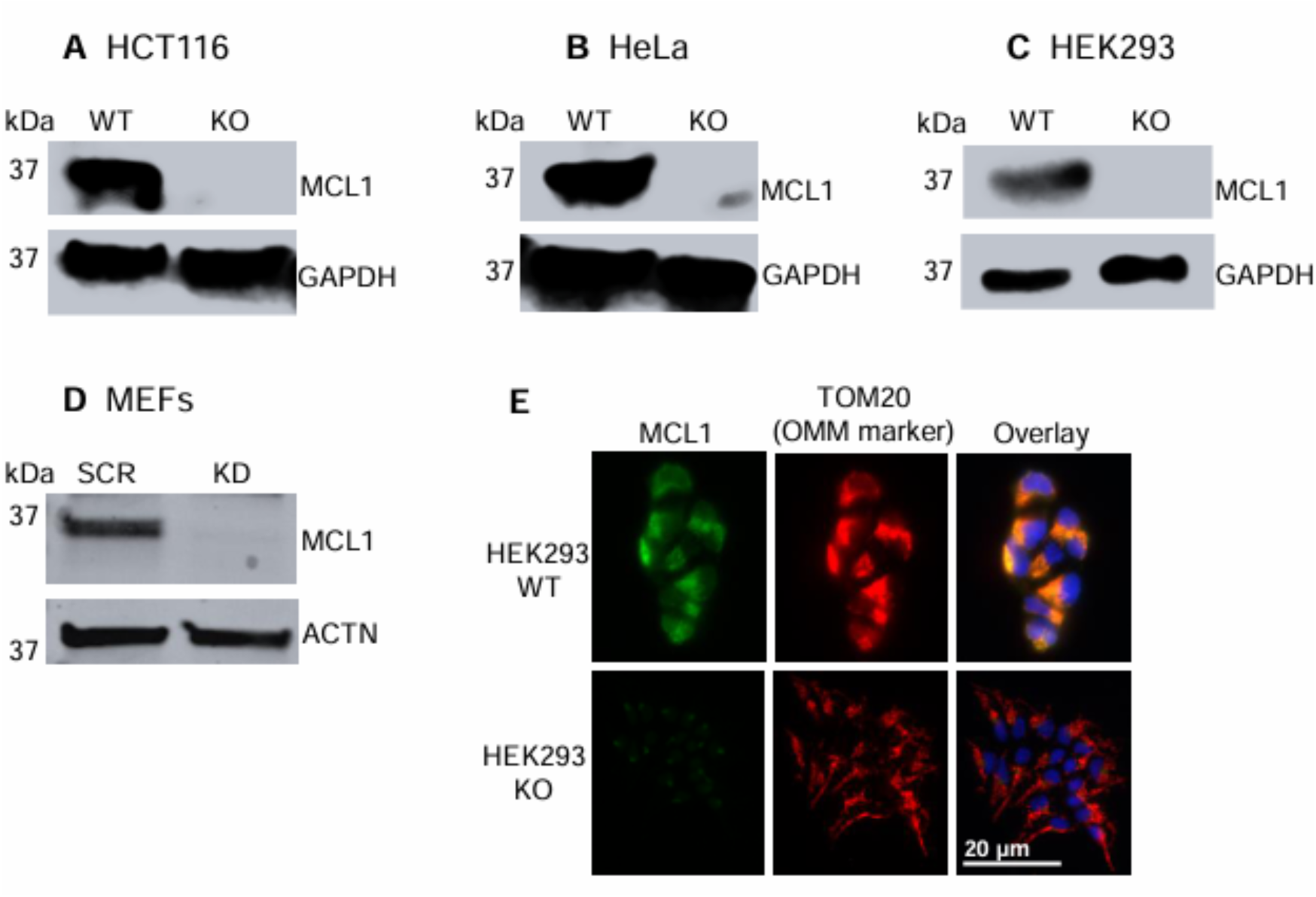
MCL1^KO/KD^ cell line and CST #5453 anti-MCL1 antibody validation by western blot in **A**). HCT116, **B**). HeLa, **C**). HEK293 and **D**). MEFs and by IF microscopy in **E**). HEK293 cells. SCR = scramble siRNA (negative control). IF images were taken at 40x magnification. Data are representative of three independent experiments.

### MCL1 loss does not increase chemosensitivity

Prior studies suggested that deleting MCL1 sensitizes cancer cell lines to chemotherapy. We repeated these assays.^47^ WT and MCL1^KO/KD^ HCT116, HeLa, and MEFs (5,000 cells/well in 96- well plates) were exposed for 48 h to paclitaxel, vincristine, vinorelbine, doxorubicin, or etoposide. Representative viability curves for HCT116, HeLa, and MEF cells are shown in **Fig. S1A-O**; pGI50 comparisons are in **Fig. 2A-C**. The same assays were also performed on WT vs. MCL1^KO^ HEK293 cells (**Fig. S1P-T**). We show in **Fig. S1U-V** that MCL1 expression remains knocked down in MEFs for the entire duration of the experiment. Drug sensitization from MCL1 removal never exceeded two-fold, and most drug-cell line combinations showed no difference.

**Figure 2.**
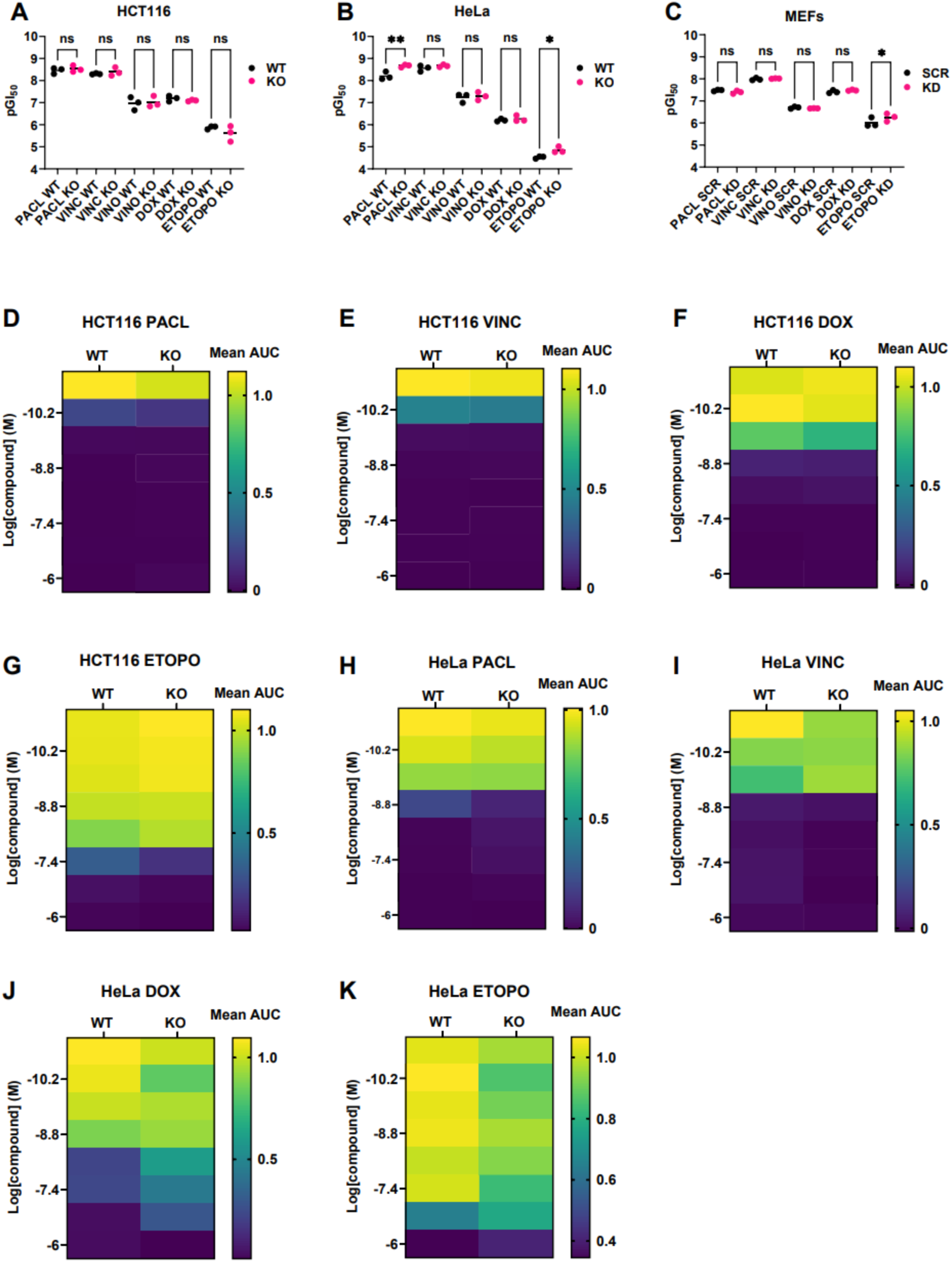
Summary of results from chemosensitivity studies on WT vs. MCL1^KO/KD^ cells in both 96-well and 384-well plate formats. One-way ANOVA with multiple comparisons of pGI50 values for each WT and MCL1^KO/KD^ pair are shown for **A)**. HCT116, **B)**. HeLa and **C)**. MEFs as a result of 48-h chemotherapy treatment in 96-well plates. Multiplicity adjusted P values were obtained for each comparison to determine statistical significance; P > 0.05 (ns), P ≤ 0.05 (*), P ≤ 0.01 (**); ns = not significant. Three pGI50 values were extracted for each WT and MCL1^KO/KD^ cell line treated with the five chemotherapies since cell viability assays were performed in biological triplicate. SCR = scramble siRNA (negative control). Heatmaps depicting the comparison of mean AUC values for WT and MCL1^KO^ **D-G)** HCT116 and **H-K)**. HeLa cell lines treated with dose titrations of PACL, VINC, VINO, DOX or ETOPO in 384-well plates. Areas were obtained under each percent confluence vs. time growth curve shown in **Figure S2A-B**. AUCs were averaged from treatments performed in technical triplicate.

To extend these results, we monitored low density cell cultures (500 cells/well in 384-well plates) by Incucyte® for 7 days (**Fig. S2A-B**). Area under the growth curve (AUC) analysis (**Fig. 2D-K**) and endpoint viability by CTG® (**Fig. S2C-J**) are revealed were quantified. We observed no difference in the growth of MCL1^KO^ and WT cells in the presence of chemotherapy. Thus, MCL1 loss does not enhance chemosensitivity in HCT116 or HeLa cells.

### Forced nuclear MCL1 does not restore chemoresistance

To test whether nuclear MCL1 promotes chemoresistance, we re-expressed 3XFLAG-tagged wild-type MCL1 (MCL1^WT^) or a construct bearing two nuclear localization sequences (MCL1^NLS^) in MCL1^KO^ HCT116 and HeLa cells. Subcellular fractionation confirmed that MCL1^NLS^ accumulates in the nucleus, while MCL1^WT^ is distributed between the nucleus and the cytosol (**Fig. 3A-C**). Neither construct altered drug responses. (**Fig. 3D-E**; **Fig. S3**).

**Figure 3.**
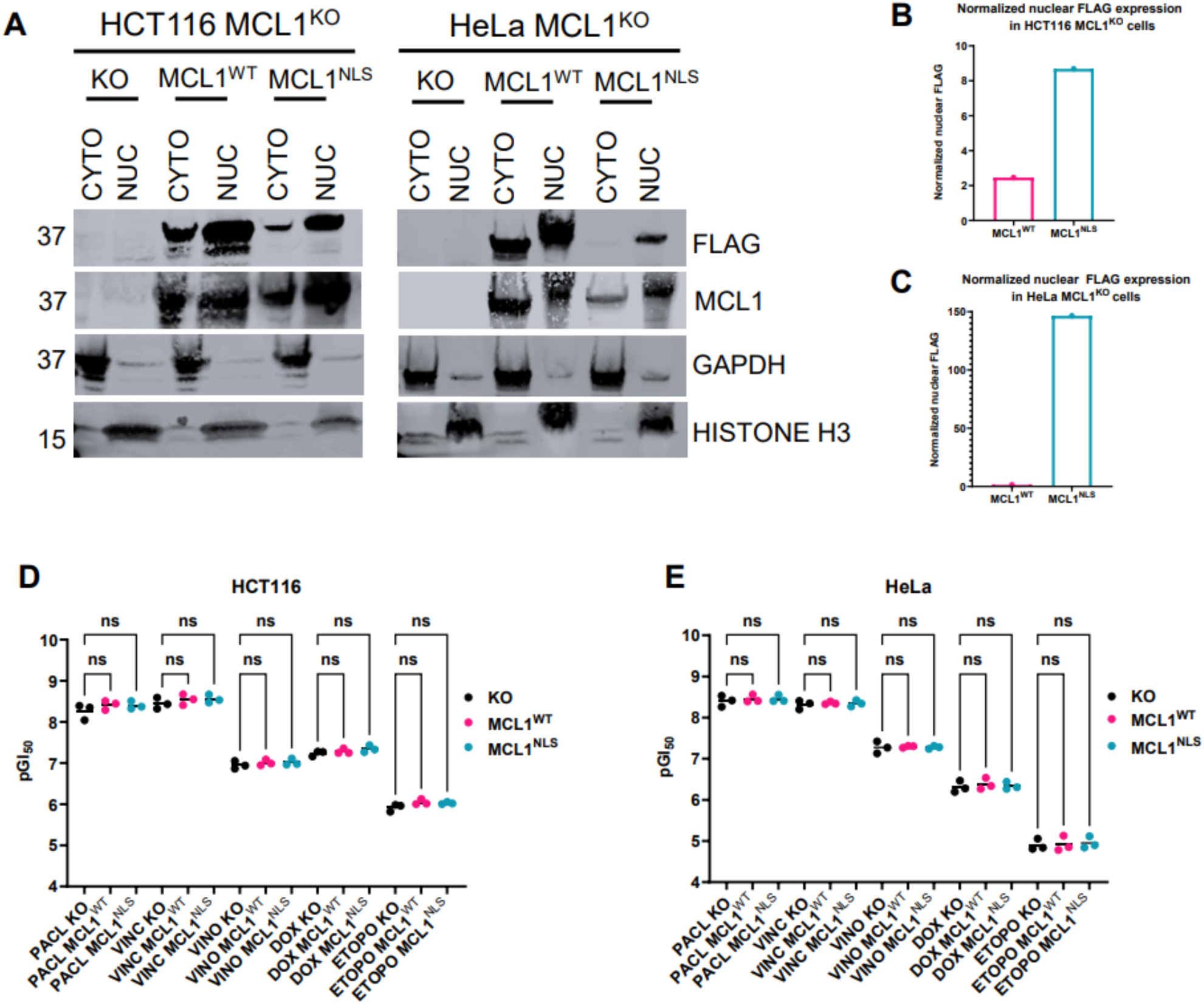
Restoration of MCL1^WT^ and overexpression of MCL1^NLS^ in MCL1^KO^ cell lines to study their effects on chemoresistance. **A)**. Validation of MCL1^WT^ and MCL1^NLS^ localization in MCL1^KO^ HCT116 and HeLa cells by subcellular fractionation. CYTO = cytosolic fraction; NUC = nuclear fraction. **B-C)**. Normalization and quantitation of nuclear FLAG expression of MCL1^WT^ and MCL1^NLS^ in MCL1^KO^ HCT116 and HeLa cells, respectively. One-way ANOVA with multiple comparisons of pGI50 values for MCL1^KO^ vs. KO cell lines reconstituted with MCL1^WT^ or MCL1^NLS^ in **D)**. HCT116 and **E)**. HeLa for each chemotherapy treatment. Multiplicity adjusted P values were obtained for each comparison to determine statistical significance; P > 0.05 (ns), P ≤ 0.05 (*), P ≤ 0.01 (**); ns = not significant. Three pGI50 values were extracted for each KO and KO cell lines reconstituted with MCL1^WT^ or MCL1^NLS^ treated with the five chemotherapies since cell viability assays were performed in biological triplicate.

### DepMap analysis supports experimental findings

Finally, to extend our observations beyond HCT116 and HeLa cells, we made use of the Broad Institute’s Cancer Dependency Map^51^ to assess the correlation between MCL1 expression and chemosensitivity across hundreds of cancer cell lines and dozens of lineages. MCL1 expression did not correlate with sensitivity to vincristine, doxorubicin, or etoposide. A weak, nonsignificant trend suggested the opposite (**Fig. S4**). Therefore, publicly available pan-cancer data agree with our results.

### MCL1 does not accumulate in the nucleus after chemotherapy

Earlier work reported treatment-induced nuclear import of MCL1 in HCT116 and HeLa cells.^34, 36, 37^ We examined the subcellular distribution of MCL1 after exposure to several drugs at published doses and times. Subcellular fractionation (**Fig. 4A-B**) and IF (**Fig. 4C, Fig. S5**) showed a consistent low-level nuclear pool of MCL1, partly of the short nuclear form (snMCL1),^33^ but we observed no treatment-dependent increase as previously suggested.^34, 36, 37, 47, 52^ Additional chemotherapeutic agents yielded the same outcome (**Fig. S5**). Thus, chemotherapy does not drive MCL1 into the nucleus.

**Figure 4.**
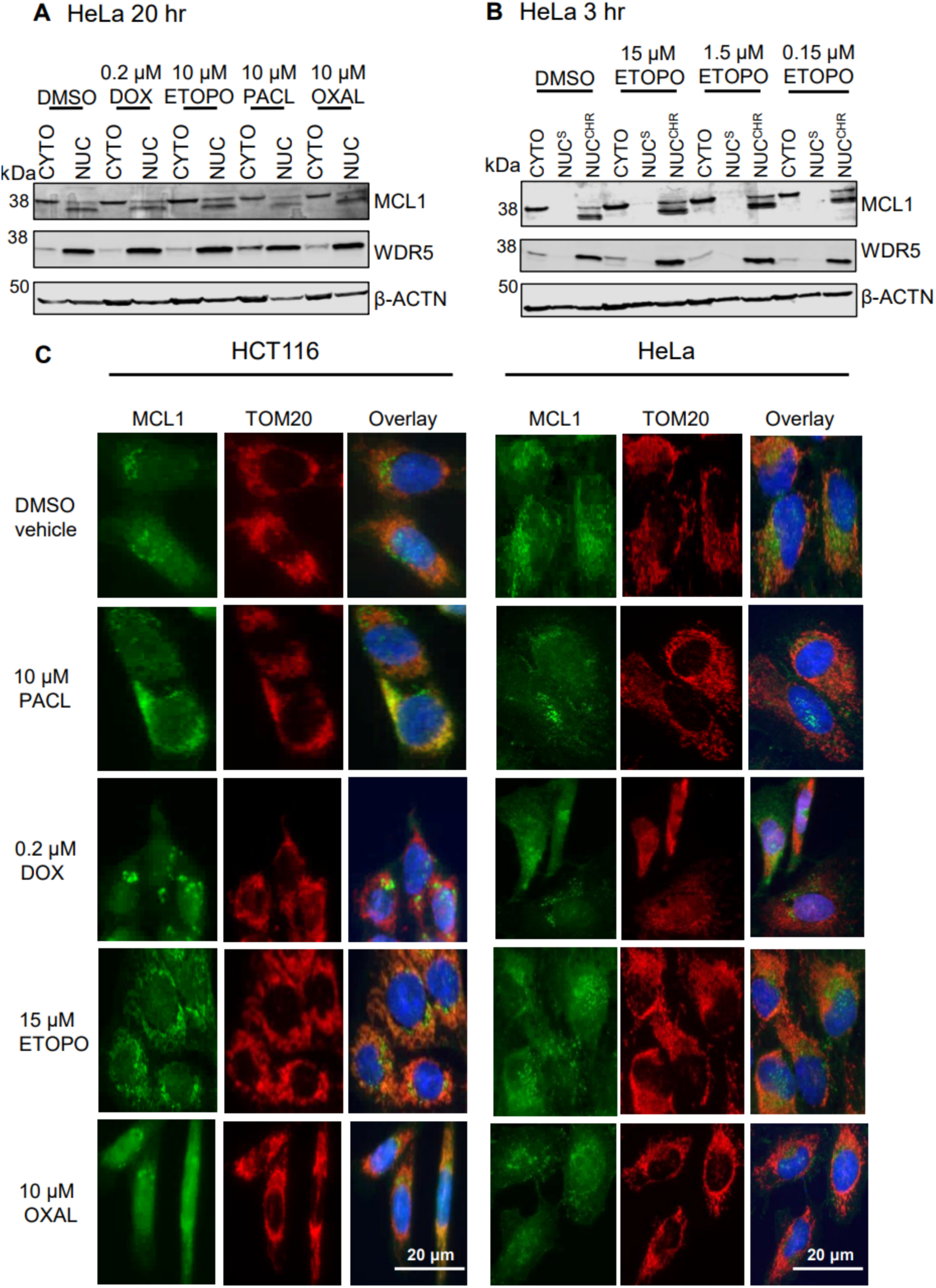
MCL1 nuclear translocation assays in HCT116 and HeLa cells. **A)**. Subcellular fractionation assessment of MCL1 nuclear translocation in response to overnight (20 hr) doxorubicin (DOX), etoposide (ETOPO), paclitaxel (PACL) and oxaliplatin (OXAL) treatments in HeLa cells. CYTO = cytosolic fraction; NUC = nuclear fraction. **B)**. Subcellular fractionation assessment of MCL1 nuclear translocation in response to 3 hr etoposide treatments. NUC^S^ = soluble nuclear proteins; NUC^CHR^ = chromatin-bound nuclear proteins. **C)**. IF evaluation of MCL1 nuclear translocation in response to 20 hr chemotherapy treatments in HCT116 and HeLa cells. IF images were taken at 40x magnification. Data are representative of two independent experiments.

### Mapping the nuclear interactome of MCL1

Previous studies, mainly by co-IP and IF colocalization, proposed nuclear partners of MCL1 such as PCNA,^43^ CDK1,^33, 53^ MCM5,^47^ CHK1,^36^ and γH2AX.^39^ Because we saw neither MCL1 nuclear translocation or MCL1-dependent chemoresistance, we used TurboID proximity biotinylation^48^ to survey nuclear interactors of MCL1 in live cells (**Fig. 5A**). To the best of our knowledge, there are no existing reports of mapping MCL1 PPIs by proximity biotinylation, which is a gold-standard method that outperforms co-IP because it can pick up transient or weak interactors of a protein in the native cellular context prior to lysis.^54^

**Figure 5.**
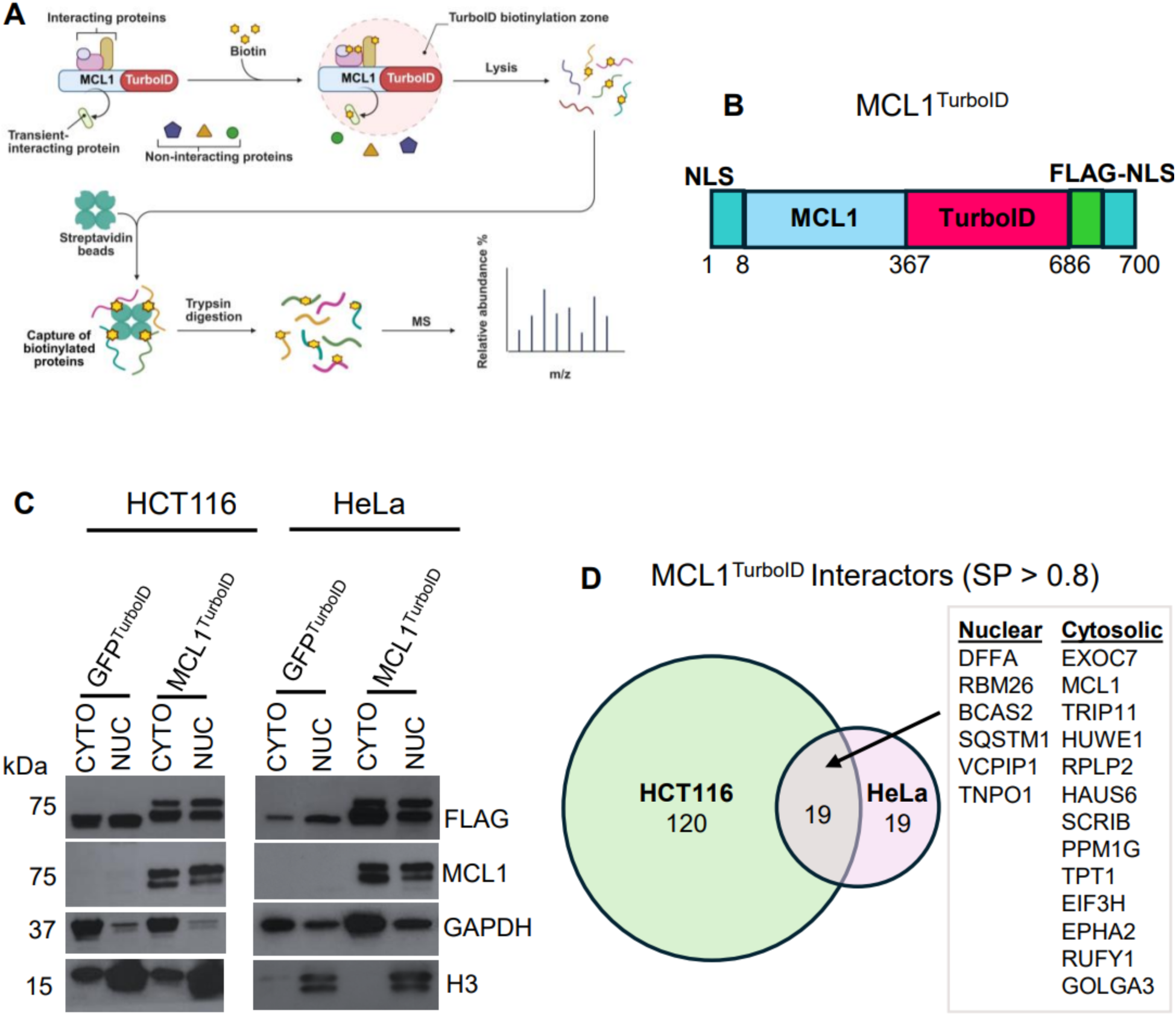
Proximity biotinylation experimental design and results. **A)**. Schematic of proximity biotinylation approach with arbitrary MCL1-TurboID fusion construct (figure modified from *BioRender*). **B).** MCL1^TurboID^ plasmid fusion construct design (75 kDa, 700 aa). **C)**. Validation of GFP^TurboID^ and MCL1^TurboID^ nuclear localization in HCT116 and HeLa cells by subcellular fractionation. CYTO = cytosolic fraction; NUC = nuclear fraction. **D)**. Venn diagram comparing number of newly defined MCL1 interactors between HCT116 and HeLa cells with SAINT SP > 0.8. Box shows the 18 shared MCL1 interactors (both nuclear and cytosolic) between both cell lines. Proximity biotinylation experiments were performed in technical triplicate.

We compared the interactome of full-length MCL1 fused to two NLS and a C-terminal TurboID (MCL1^TurboID^; **Fig. 5B**) to an identical GFP^TurboID^ construct. Subcellular fractionation verified the presence of both fusion proteins in the nucleus (**Fig. 5C**).

We identified 120 unique MCL1 interactors in HCT116 cells, 19 unique MCL1 interactors in HeLa cells and 18 common interactors in both cell lines (**Files S1-3**; **Fig. 5D**; **Fig. S6**). We identified 16 novel interactors, including six nuclear-localized proteins: DNA fragmentation factor alpha (DFFA), RNA binding protein 26 (RBM26), pre-mRNA-splicing factor SPF27 (BCAS2), sequestosome-1 (SQSTM1), the deubiquitinating enzyme VCPIP1, and the nuclear-pore component transportin-1 (TNPO1). None of the previously reported nuclear interactors of MCL1 were identified with a probability SP > 0.8, although we confidently identified MCL1 itself, MCL1’s known partner E3 ligase Mule aka HUWE1,^55, 56^ and one known cytoplasmic MCL1 interactor (translationally-controlled tumor protein TPT1 aka p23).^57, 58^ In aggregate, our data do not support previous reports on the specific interaction of nuclear MCL1 with DNA damage or cell cycle- regulating proteins in cancer cells.

## DISCUSSION

MCL1’s established anti-apoptotic activity is mitochondrial, yet its reported non-canonical roles in development, hematopoiesis, and therapy response make it a unique BCL2 family member. Elucidating these non-apoptotic functions may uncover drug targets with wider therapeutic windows than BH3-groove inhibitors–for example, by blocking MCL1’s nuclear interactions. Several studies have proposed that MCL1 translocates to the nucleus after chemotherapy, thereby fostering “chemoresistance,” often inferred from surrogate read-outs such as beta-galactosidase activity or γH2AX.^37, 41, 42, 59^ Direct effects on drug-induced cell death, however, have not been examined systematically.

Taken together, our data do not support a chemoprotective role for MCL1. MCL1 deletion or knockdown failed to sensitize HCT116, HeLa, or MEF cells to five chemotherapies, and forced re-expression of MCL1–including a nucleus-restricted variant–did not restore chemoresistance to cells lacking MCL1. Moreover, subcellular fractionation and IF showed that chemotherapy does not increase nuclear MCL1. One potential reason for this discrepancy is our use of a rigorously validated anti-MCL1 monoclonal antibody rather than the discontinued polyclonal antibody cited previously. TurboID proximity biotinylation revealed previously unreported nuclear interactors but no partners involved in the DNA damage response or cell cycle regulation. Thus, while a constitutive nuclear pool of MCL1 exists and engages distinct proteins, the present evidence argues against a role for chemoresistance. Further work should focus on defining the biological consequence of these newly identified interactors, independent of drug response.

## AUTHOR CONTRBUTIONS

**K.A. Emhoff**: Resources, data curation, formal analysis, validation, investigation, visualization, writing-original draft, writing-review and editing. **K. Chung**: Visualization, methodology, writing-review. **D. Zhang**: Resources, data curation, formal analysis. **B. Willard**: Resources, data curation, formal analysis. **T. Chan**: Resources. **B.K. Jha**: Resources. **S.R. Stauffer**: Resources, investigation. **J.A. Coker**: Conceptualization, resources, data curation, formal analysis, investigation, methodology, supervision, writing-original draft, writing-review and editing. **J.J. Melenhorst**: Conceptualization, supervision, resources, investigation, project administration, writing-original draft, writing-review and editing.

## ACKNOWLEDGEMENTS

This work was supported by NIH grant R01CA244958 (to J.J.M). We thank Cleveland Clinic Research’s Imaging Core for the use of their microscope in obtaining IF images, and the Cell Services Core for providing cell lines and culture media used in this work.

