## Supplemental Information for "MCL1 may not mediate chemoresistance"

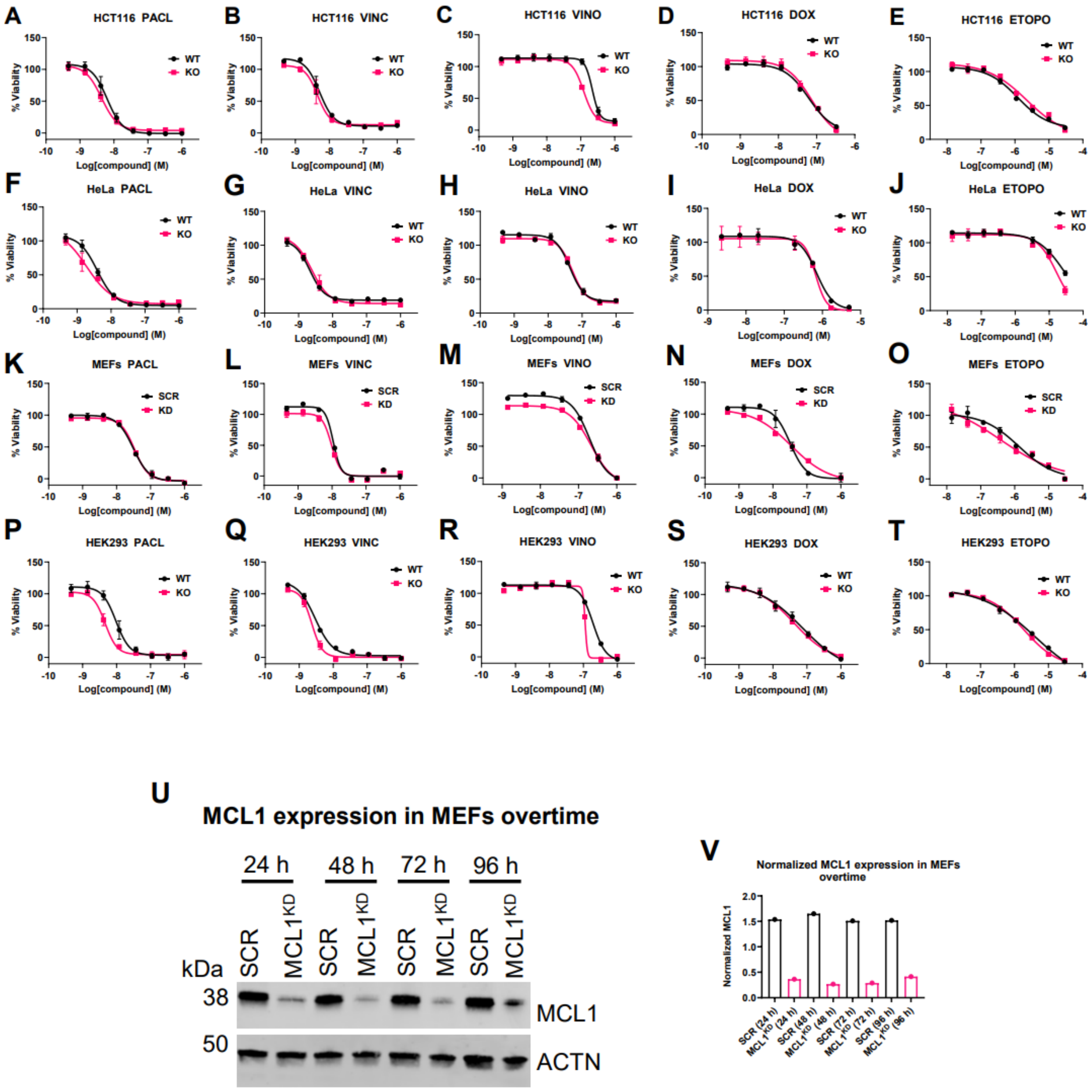


**Figure S1**. Chemosensitization studies on WT vs. MCL1$^{KO/KD}$ cell lines by CTG®. Cell viability was quantified with CTG® after 48 h treatment with paclitaxel (PACL), vincristine (VINC), vinorelbine (VINO), doxorubicin (DOX), or etoposide (ETOPO) in **A-E)**. HCT116, **F-J)**. HeLa, **K-O)**. MEF, and **P-T)**. HEK293 cells. Assays were performed in biological triplicate, with each one in technical duplicate. All data points on viability curves represent average % viability ± SD. **U-V).** MCL1 expression monitored over 4 days in MEFs after MCL1 siRNA transfection; data are representative of two independent experiments. SCR = scramble siRNA (negative control).

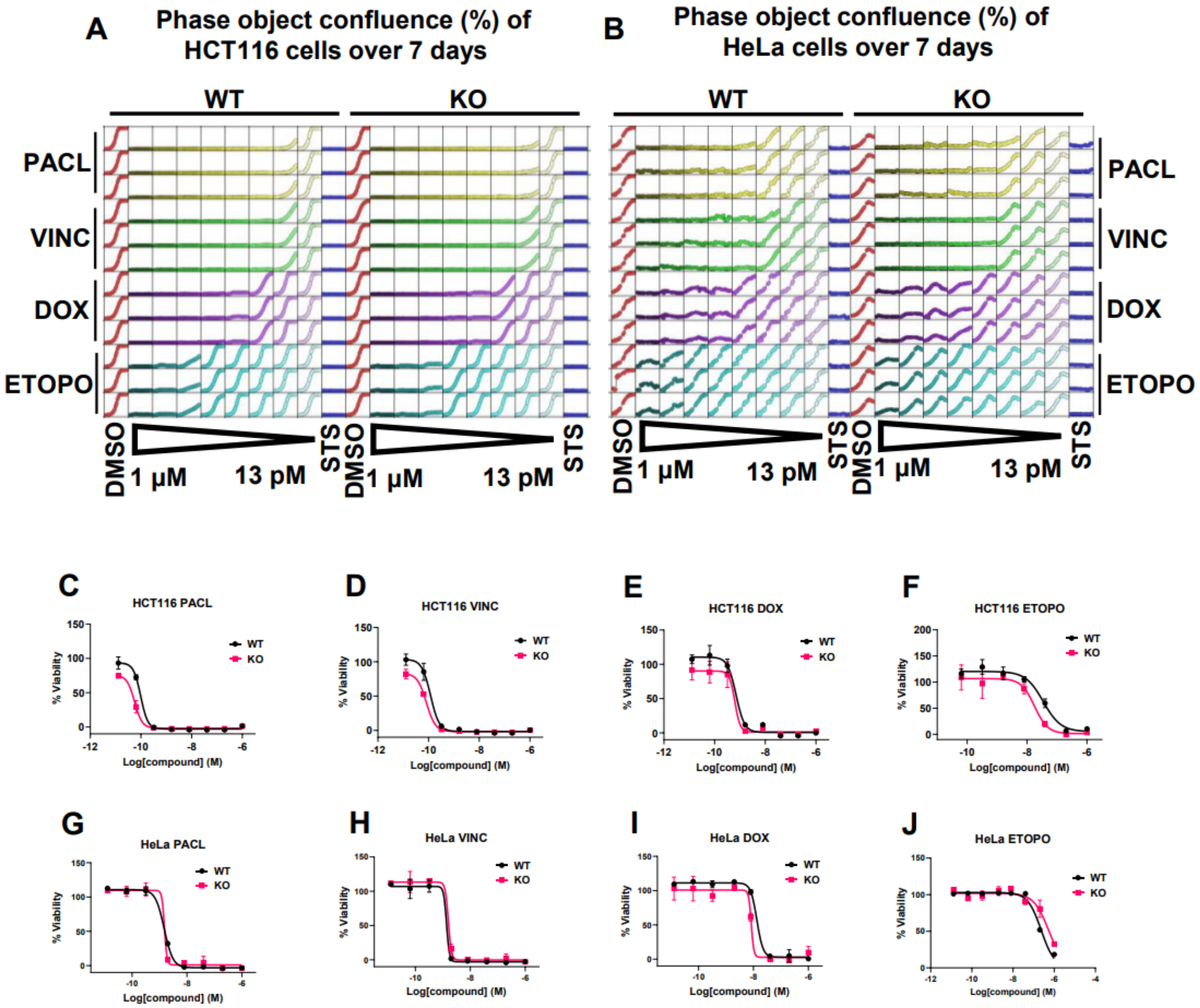


**Figure S2**. Live-cell analysis of phase object confluence (%) in chemotherapy-treated **A)**. WT vs. KO HCT116 and **B)**. HeLa cells on Incucyte® over 7-day period. Cell viability measured by CTG® after 7-day treatment with PACL, VINC, DOX or ETOPO in **C-F)**. HCT116 and **G-J)**. HeLa cells. Experiments **A-J** were performed in biological duplicate, with each one in technical triplicate. All data points on viability curves represent average % viability ± SD.

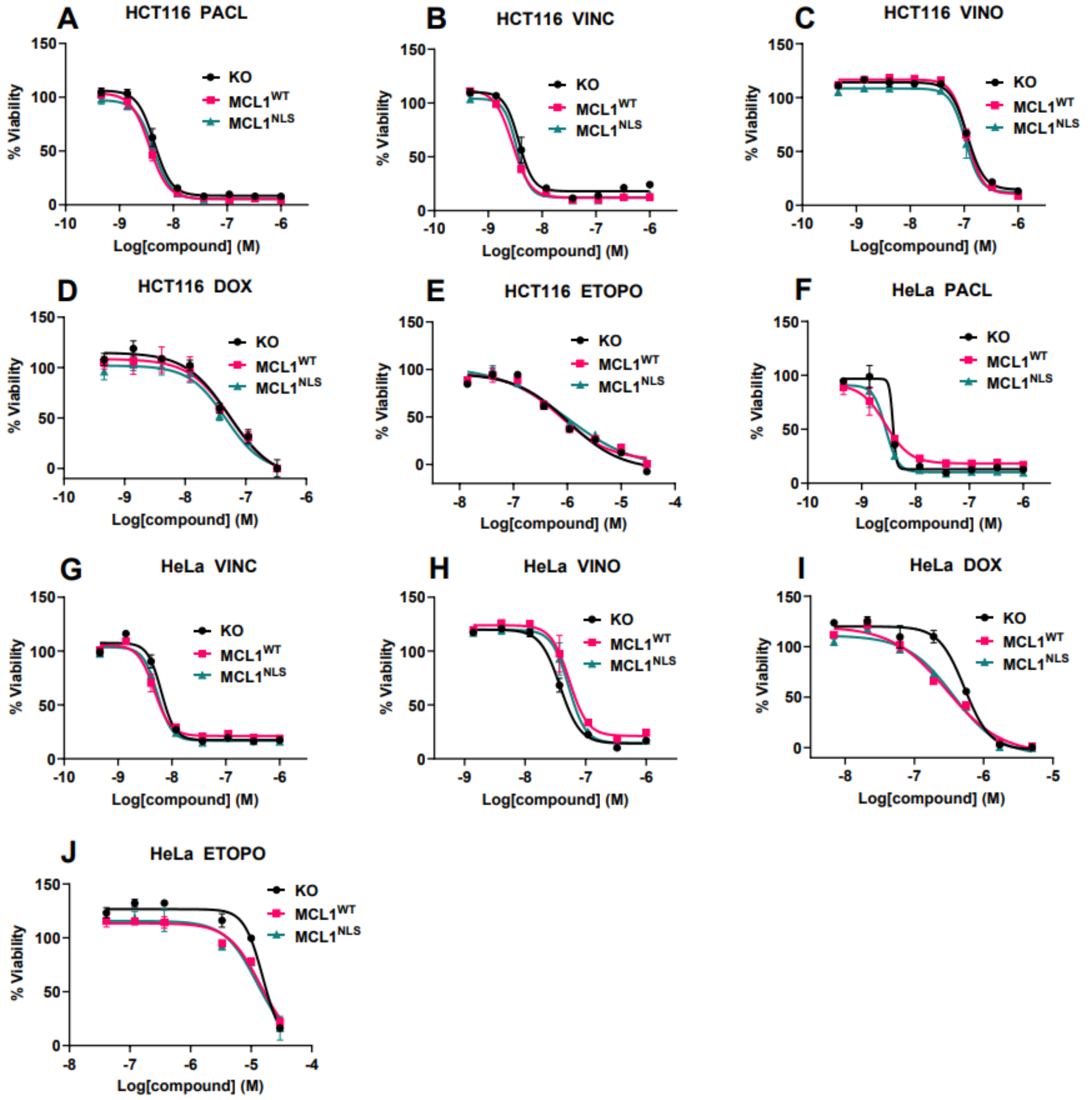


**Figure S3**. Cell viability assays on parental MCL1$^{KO}$ vs. KO cell lines reconstituted with MCL1$^{WT}$ or MCL1$^{NLS}$. Cell viability was measured by CTG® after 48 h treatment with PACL, VINC, VINO, DOX or ETOPO in **A-E)**. HCT116 and **F-J)**. HeLa cells. Cell viability assays were performed in biological triplicate, with each one in technical duplicate, so data points represent average % viability ± SD.

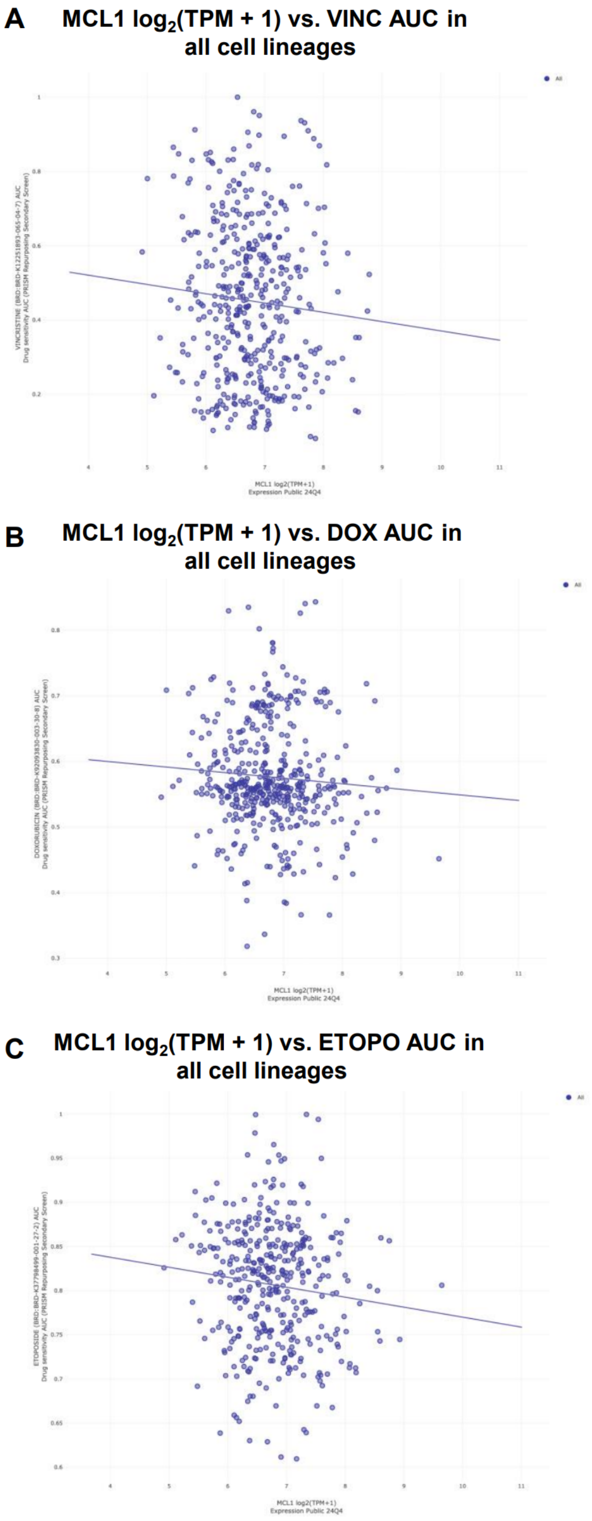


**Figure S4**. MCL1 RNA expression (log2(TPM + 1)) vs. AUC from viability experiments utilizing chemotherapeutic treatments. Graphs were generated using the publicly available datasets in the Broad Institute's DepMap and show the correlation between MCL1 expression and AUC in ~900 cancer cell lines in response to **A-B)**. VINC, **C-D)**. DOX or **E-F)**. ETOPO treatments. An AUC value of 1.0 represents 100% cell viability, whereas a value of 0.0 represents 0% cell viability.

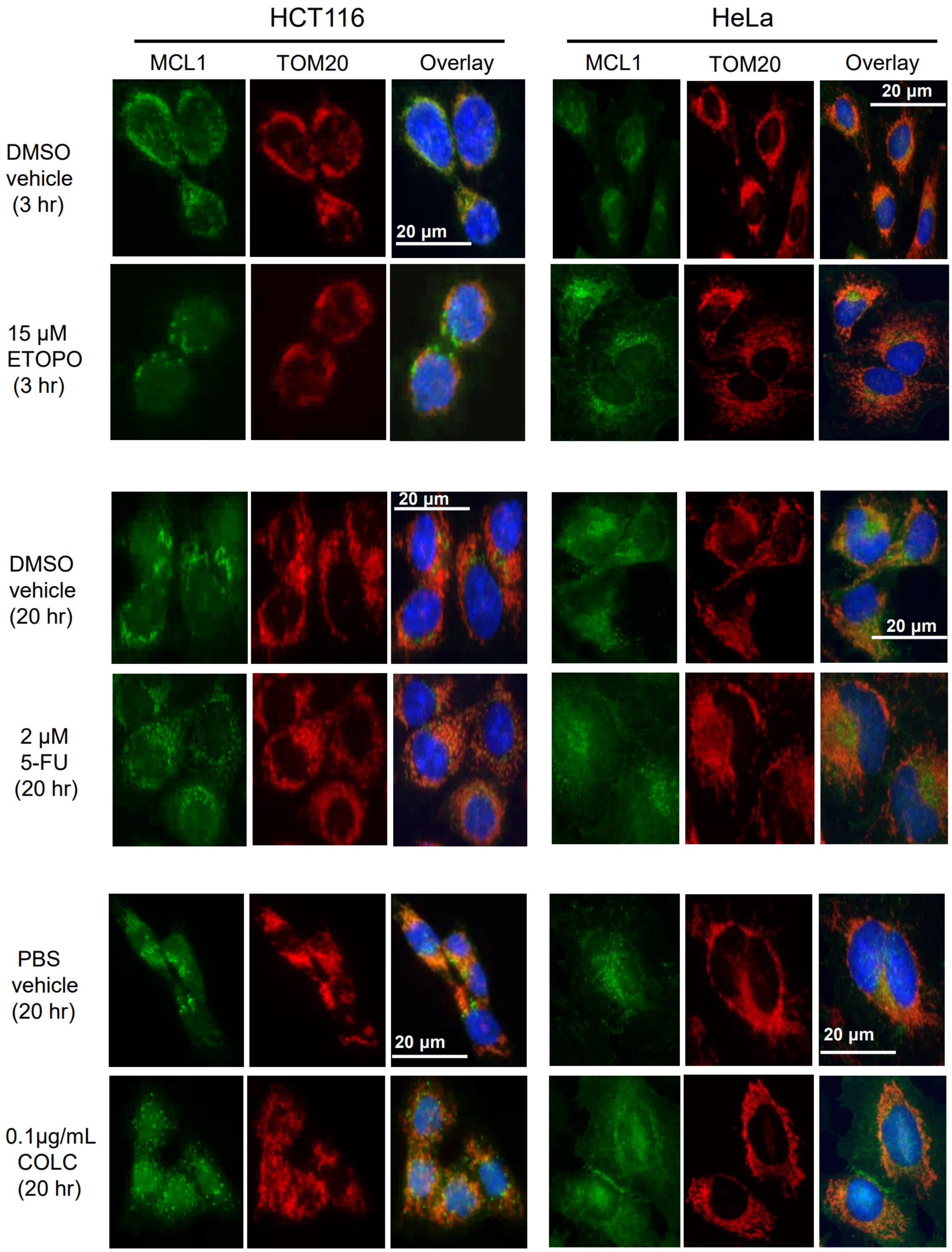


**Figure S5**. IF evaluation of MCL1 nuclear translocation in response to 3 hr 15 µM etoposide, 20 hr 2 µM 5-fluorouracil (5-FU) or 0.1 µg/mL colcemid (COLC) treatments in HCT116 and HeLa cells. IF images were taken at 40x magnification. Data are representative of two independent experiments.

**A** **MCL1$^{TurboID}$ vs. GFP$^{TurboID}$ HCT116 Cells**

SP > 0.8

*MCL1 (SP = 1.0, FC = 488 excluded from figure)*

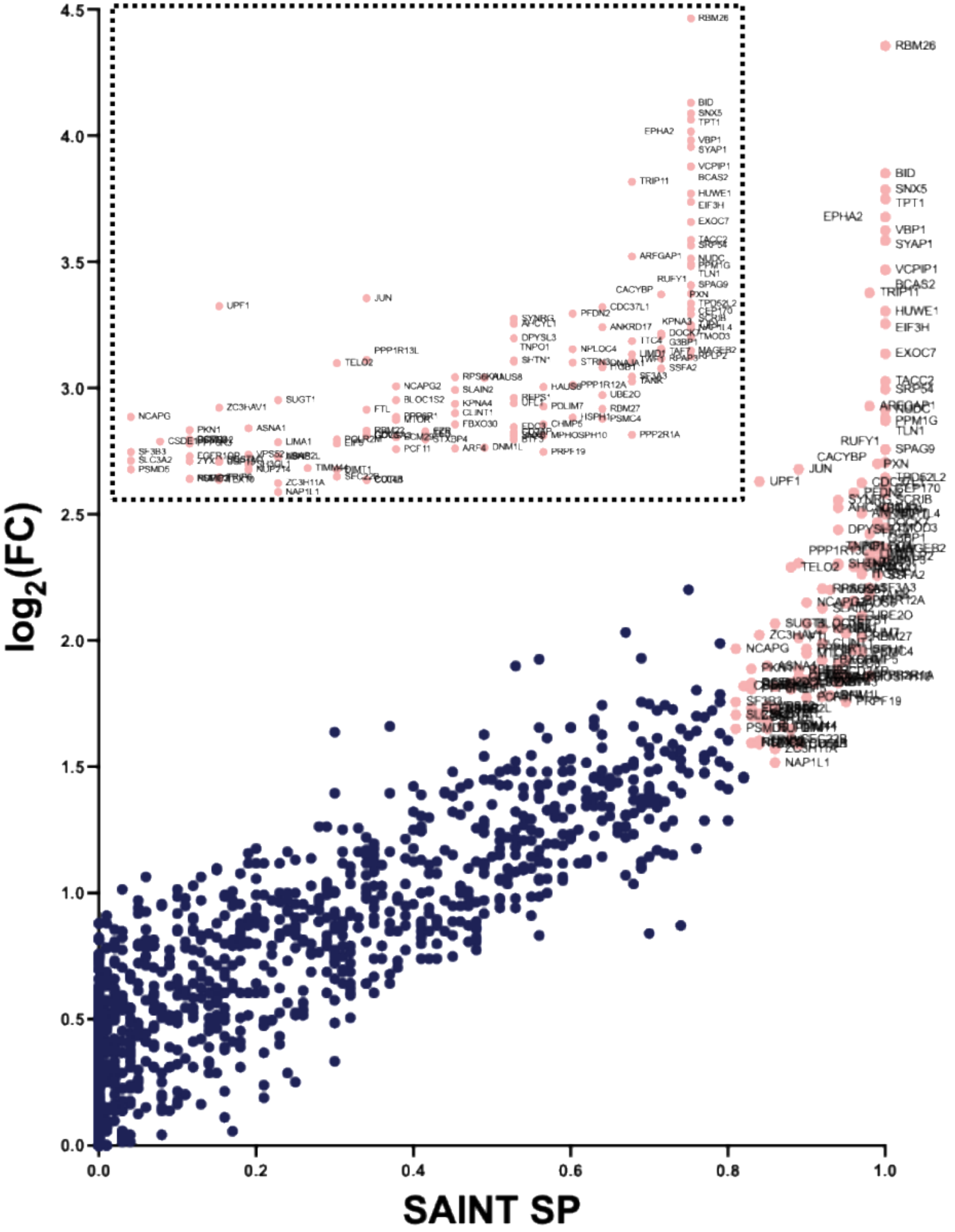

**B**

**MCL1$^{TurboID}$ vs. GFP$^{TurboID}$ HeLa Cells**

SP > 0.8

*MCL1 (SP = 1.0, FC = 335 excluded from figure)*

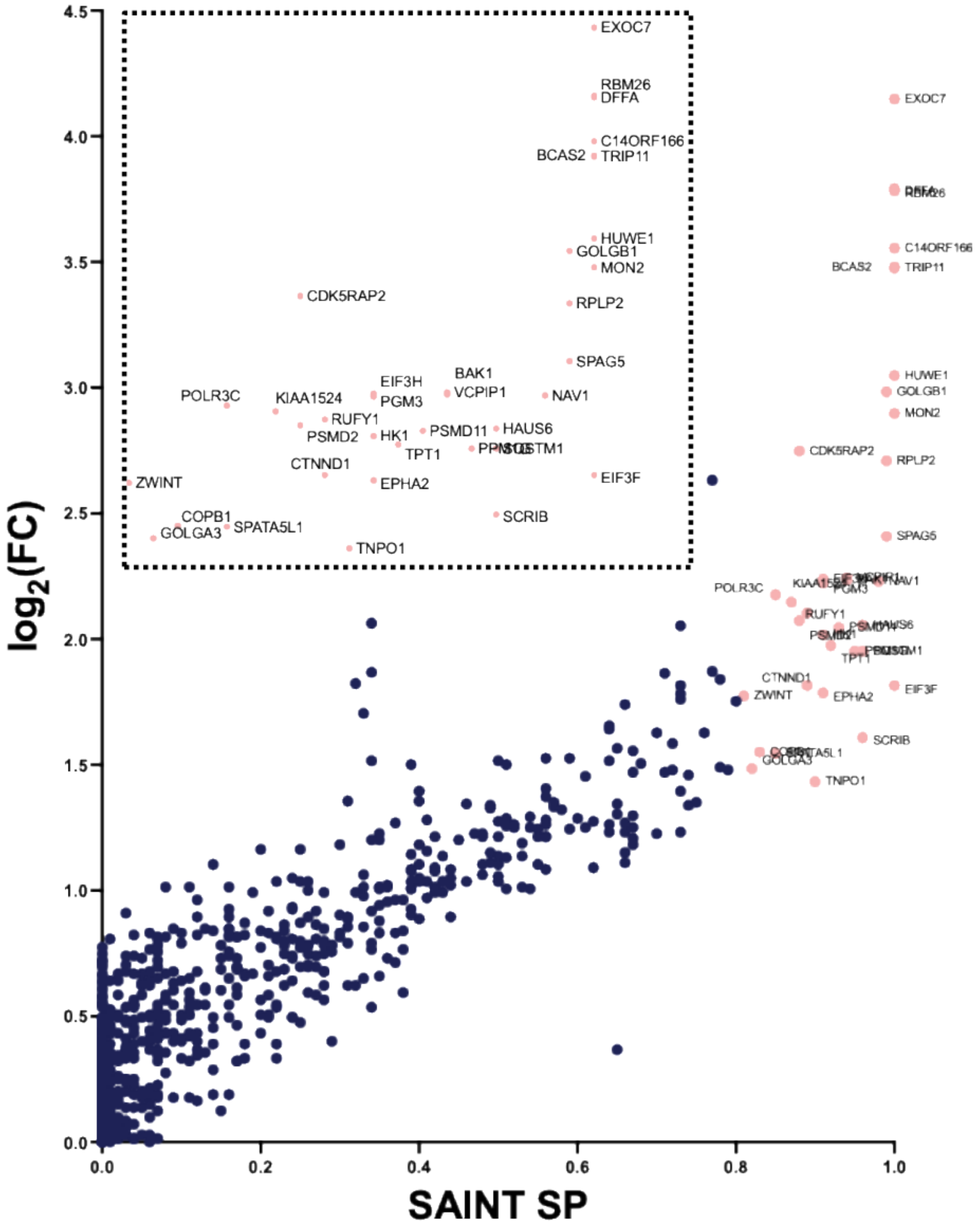

**Figure S6**. Volcano plots of MCL1 interactors enriched in $MCL1^{WT\text{-}TurboID}$ vs. $GFP^{TurboID}$ with SAINT SP > 0.8 in **A)**. HCT116 and **B)**. HeLa cells. Data set shows average SP scores from three replicates.
